# Attenuated directed exploration during reinforcement learning in gambling disorder

**DOI:** 10.1101/823583

**Authors:** A. Wiehler, K. Chakroun, J. Peters

## Abstract

Gambling disorder is a behavioral addiction associated with impairments in value-based decision-making and behavioral flexibility and might be linked to changes in the dopamine system. Maximizing long-term rewards requires a flexible trade-off between the exploitation of known options and the exploration of novel options for information gain. This exploration-exploitation trade-off is thought to depend on dopamine neurotransmission. We hypothesized that human gamblers would show a reduction in directed (uncertainty-based) exploration, accompanied by changes in brain activity in a fronto-parietal exploration-related network.

Twenty-three frequent, non-treatment seeking gamblers and twenty-three healthy matched controls (all male) performed a four-armed bandit task during functional magnetic resonance-imaging. Computational modeling using hierarchical Bayesian parameter estimation revealed signatures of directed exploration, random exploration, and perseveration in both groups. Gamblers showed a reduction in directed exploration, whereas random exploration and perseveration were similar between groups.

Neuroimaging revealed no evidence for group differences in neural representations of basic task variables (expected value, prediction errors). Our hypothesis of reduced frontal pole recruitment in gamblers was not supported. Exploratory analyses revealed that during directed exploration, gamblers showed reduced parietal cortex and substantia-nigra / ventral-tegmental-area activity. Cross-validated classification analyses revealed that connectivity in an exploration-related network was predictive of group status, suggesting that connectivity patterns might be more predictive of problem gambling than univariate effects.

Findings reveal specific reductions in strategic exploration gamblers that might be linked to altered processing in a fronto-parietal network and/or changes in dopamine neurotransmission implicated in gambling disorder.

**Significance statement:** Wiehler et al. report that gamblers rely less on the strategic exploration of unknown, but potentially better rewards during reward learning. This is reflected in a related network of brain activity. Parameters of this network can be used to predict the presence of problem gambling behavior in participants.

## Introduction

Gambling disorder (GD) has a lifetime prevalence of around 1% (Kessler et al., 2008; Lorains et al., 2011). In the DSM-5, it is classified in the category of substance use and addictive disorders, reflecting the considerable overlap in behavioral and neural correlates with substance-based addictions (Goudriaan et al., 2019). For example, activity in reward-related brain regions, including the ventral striatum and medial prefrontal cortex, has repeatedly been found to differ between healthy controls and participants with GD (Balodis et al., 2012; Leyton and Vezina, 2012; Miedl et al., 2012), though with inconsistent directionality (Clark et al., 2019).

In addition to increased temporal discounting and risk-taking (Wiehler and Peters, 2015), gamblers also exhibit cognitive impairments reflected in reduced behavioral flexibility. This includes impaired performance in the Stroop task and increased perseveration following rule changes in the Wisconsin Card Sorting Task (van Timmeren et al., 2018). State-dependent modulations of risk-attitude have been found impaired in problem gambling (Fujimoto et al., 2017). Similar impairments are observed in reversal learning, where gamblers make more perseveration errors following contingency reversals (de Ruiter et al., 2009; Boog et al., 2014), an effect that has been linked to maladaptive control beliefs about gambling outcomes, which might interfere with decision-making (Lim et al., 2015).

More generally, reward-learning entails a trade-off between exploitation of options with known value, and exploration of novel options for information gain (Wilson et al., 2021). One of the most widely used tasks to examine exploration behavior is the multi-armed-bandit task (Daw et al., 2006). Here, participants make repeated choices between multiple choice options (“bandits”) to obtain rewards. Exploitation involves tracking each bandit’s expected value and choosing the best. In contrast, exploration can be undirected due to stochastic selection of bandits (“random exploration”) (Daw et al., 2006; Schulz and Gershman, 2019). Additionally, exploration might entail a goal-directed component and depend on the bandit’s estimated uncertainty (“directed exploration”) (Speekenbrink and Konstantinidis, 2015; Schulz and Gershman, 2019; Chakroun et al., 2020).

A bilateral fronto-parietal network supports exploration, including intra-parietal sulcus and frontopolar cortex (Daw et al., 2006; Raja Beharelle et al., 2015; Chakroun et al., 2020). Although initially characterized in the context of random exploration (Daw et al., 2006), fronto-polar cortex may more specifically support directed exploration (Boorman et al., 2009, 2011; Badre et al., 2012; Zajkowski et al., 2017).

There is substantial evidence implicating the neurotransmitter dopamine (DA) in the pathophysiology of gambling disorder (Kayser, 2019). Likewise, a contribution of DA to the regulation of the exploration-exploitation trade-off is suggested both by theory (Beeler, 2012) and empirical data (Frank et al., 2009; Kayser et al., 2014; Gershman and Tzovaras, 2018; Cinotti et al., 2019; Chakroun et al., 2020). The most prominent empirical observation implicating DA in gambling comes from patients suffering from Parkinson’s disease, where higher rates of problem gambling behavior haven been liked to pharmacological DA replacement therapy (Driver-Dunckley et al., 2003; Voon et al., 2006). Gamblers may also exhibit increased pre-synaptic striatal DA levels (Boileau et al., 2014; van Holst et al., 2017), although this is controversially discussed (Majuri et al., 2017; Potenza, 2018).

We have recently shown that an elevation of DA levels via L-Dopa attenuates directed exploration in healthy controls (Chakroun et al., 2020). If one conceptualizes gambling disorder as a hyperdopaminergic state (Boileau et al., 2014; van Holst et al., 2017), this entails the prediction that gambling disorder might likewise be associated with reduced directed exploration. This hypothesis also resonates with the discussed impairments in behavioral flexibility in gambling disorder. In line with the critical role of frontal pole regions (Daw et al., 2006; Raja Beharelle et al., 2015; Zajkowski et al., 2017) and prefrontal dopamine (Frank et al., 2009) in exploration, we further hypothesized that reduced frontal pole recruitment might contribute to reduced exploration in gambling disorder. We addressed these hypotheses in a group of frequent gamblers (with sixteen out of twenty-three meeting the diagnostic criteria for gambling disorder) and healthy matched controls using an established 4-armed bandit task during functional magnetic resonance imaging (fMRI, Daw et al., 2006).

## Materials and Methods

### Sample

We investigated a sample of n=23 frequent gamblers (age mean [SD]=25.91 [6.47], all male). Sixteen gamblers fulfilled four or more DSM-5 criteria of gambling addiction (mean [SD]=6.31 [1.45], previously defined as pathological gamblers). Seven gamblers fulfilled one to three criteria (mean [SD]=2.43 [0.77], previously defined as problem gamblers). All participants reported no other addiction except for nicotine. Current drug abstinence was verified via urine drug screening. All participants reported no history of other psychiatric or neurological diagnoses except depression. No participant was undergoing any psychiatric treatment. Current psychopathology was controlled using the Symptom Checklist 90 Revised (SCL-90-R) questionnaire (Schmitz et al., 2000) and depression symptoms were assessed via the Beck Depression Inventory-II (BDI-II, Osman et al., 2004). To characterize gambling behavior, we conducted the German gambling questionnaire “Kurzfragebogen zum Glücksspielverhalten” (KFG, Petry, 1996), the German version of the South Oaks Gambling Screen (SOGS, Lesieur and Blume, 1987) and the Gambling Related Cognitions Scale (GRCS, Raylu and Oei, 2004). Participants were recruited via advertisements placed on local internet boards but were not searching for treatment.

We recruited n=23 healthy control participants, matched for age, gender, education, income, alcohol (Alcohol Use Disorders Identification Test, AUDIT, Saunders et al., 1993) and nicotine consumption (Fagerstrom Test of Nicotine Dependence, FTND, Heatherton et al., 1991), see **Table 1.** Four of these control participants were included from an earlier study that used the exact same task and imaging protocol (Chakroun et al., 2020). To rule out drug or order effects, we included the first imaging session of participants who completed the placebo condition first. Furthermore, these four participants were selected to maximize matching to the gamblers group in terms of age, education and income. All results were significant without these four additional participants.

**Table 1.** Summary of demographics and group matching statistics. FTND: Fagerstrom Test of Nicotine Dependence, AUDIT: Alcohol Use Disorders Identification Test, KFG: Kurzfragebogen zum Glücksspielverhalten, SOGS: South Oaks gambling screen, BDI-II: Beck Depression Inventory-II. GD: Gambling disorder. HC: Healthy controls.

|  | Gamblers (n=23) |  | Controls (n=23) |  |  |  |  |
| --- | --- | --- | --- | --- | --- | --- | --- |
|  | Mean | SD | Mean | SD | t | df | p |
| Age | 25.91 | 6.47 | 26.52 | 5.92 | -0.33 | 43.65 | 0.74 |
| School years | 11.64 | 1.77 | 11.91 | 1.35 | -0.60 | 41.04 | 0.55 |
| Monthly income in EUR | 1439.86 | 835.84 | 1093.76 | 460.07 | 1.72 | 34.81 | 0.09 |
| FTND | 2.14 | 2.58 | 1.83 | 2.15 | 0.44 | 42.58 | 0.66 |
| AUDIT | 6.09 | 7.14 | 6.52 | 4.57 | -0.24 | 37.44 | 0.81 |
| DSM-5 score | 5.13 | 2.22 | 0.09 | 0.29 | 10.80 | 22.74 | <0.01 |
| KFG | 25.91 | 14.15 | 0.48 | 1.12 | 8.59 | 22.28 | <0.01 |
| SOGS | 8.64 | 4.46 | 0.22 | 0.52 | 9.00 | 22.60 | <0.01 |
| BDI-II | 15.41 | 11.41 | 7.61 | 7.94 | 2.69 | 39.27 | 0.01 |

All participants provided informed written consent before participation and the study procedure was approved by the local institutional review board (Hamburg Board of Physicians).

### Task and Procedure

Participants completed two sessions of testing on separate days. The first session included all questionnaires and an assessment of the spontaneous eye-blink rate, that was published previously (Mathar et al., 2018). The second session started with a training session of the task, followed by functional and structural magnetic resonance imaging (MRI). Subsequently, they performed an additional task in the MRI that will be reported elsewhere.

We used a previously described 4-armed bandit task (Daw et al., 2006). We applied the same task as in the original publication, with the exception that we replaced slot machine images for each bandit with colored boxes (see **Figure 1 A**). On each trial, participants selected one of four bandits. They received a payout between 0 and 100 points for the chosen bandit, which was added to a total score. The points that could be won on each trial were determined by Gaussian random walks, leading to payouts fluctuating slowly throughout the experiment (see **Figure 1 B**, for mathematical details see below). Participants completed 300 trials in total that were split into four blocks separated by short breaks. We instructed participants to gain as many points as possible during the experiment. Reimbursement was a fixed baseline amount plus a bonus that depended on the number of points won in the bandit task. In total, participants received between 70 and 100 Euros for participation.

**Figure 1.**
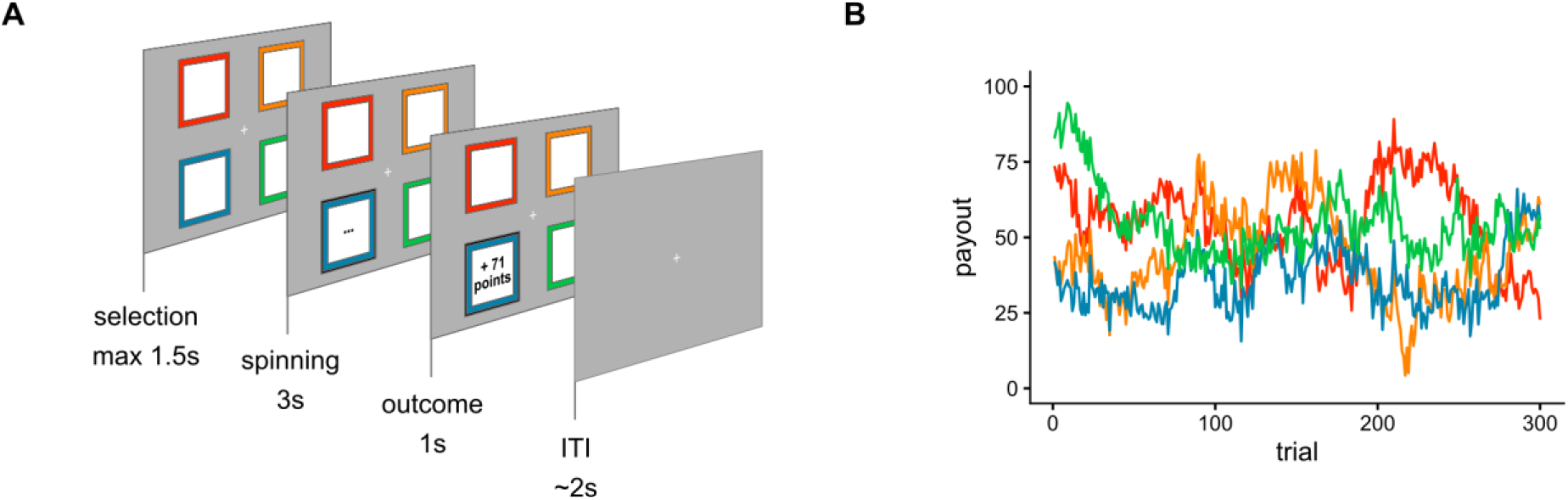
Task. A: One trial of the bandit task. On each trial, participants choose between four bandits on the screen and received a payout in reward points. **B**: Payouts fluctuated across the 300 trials of the experiment according to Gaussian random walks. Here, one example set of random walks is shown. Colors correspond to bandits in A.

### Computational modeling

To quantify exploration behavior, participants’ choices were fitted with several reinforcement learning models of varying complexity. We first implemented a Q-learning model (Sutton and Barto, 1998). Here, participants update the expected value (Q-value) of the *i*th bandit on trial *t* via a prediction error *δ_t_*:

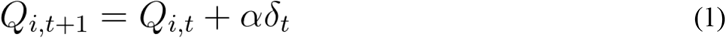

with

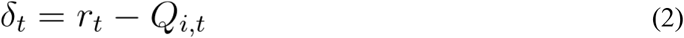

Here, *Q* is the expected value of the *i*th bandit on trial *t*, *α* is a constant learning rate, that determines the proportion of the prediction error *δ_t_* that is used for the value update, and *r_t_* is the reward outcome on trial *t*. In this model, unchosen bandits are not updated but retain their previous Q-values.

Q-values are transformed into action probabilities, using a softmax choice rule:

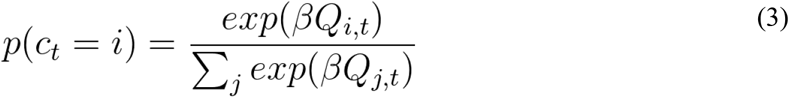

Here, *p* is the probability of choice *c_t_* of bandit *i* in trial *t*, given the estimated values *Q* from equation 1 for all *j* bandits. *β* denotes an inverse temperature parameter, that models choice stochasticity: For greater values of *β,* choices become more dependent on the learned Q-values. Conversely, as *β* approaches 0, choices become more random. In this model, *β* controls the exploration-exploitation trade-off such that for higher values of *β*, exploitation dominates, whereas exploration increases as *β* approaches 0. Note, however, that this model does not incorporate uncertainty about Q-values, as only mean Q-values are tracked.

We next examined a Bayesian learner model (Kalman filter) that was also applied by Daw et al. (2006). This model assumes that participants use a representation of the Gaussian random walks that constitute the task’s payout structure. Thus, irrespective of the choice, mean and variance of each bandit *i* are updated on each trial *t* as follows:

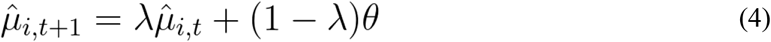

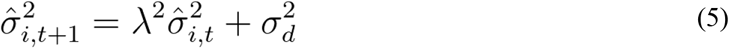

Here, *μ* is the mean expected value, *σ* is the standard deviation of the expected value, *λ* is a decay rate (fixed to 0.9836), *θ* is the decay center (fixed to 50), and *σ_d_* is the standard deviation of the diffusion noise (fixed to 2.8). Note that these equations are used to generate the Gaussian walks (see Daw et al. 2006)). That is, without sampling, each bandits’ mean value slowly decayed towards *θ*, and standard deviations increased *σ_d_* units per trial.

The bandit chosen on trial *t* (*c_t_)* is additionally updated using a delta rule similar to equation (2):

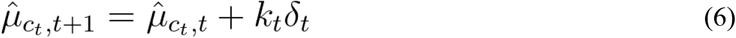

with

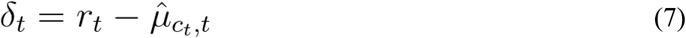

and

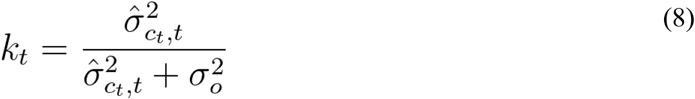

Equation 6 is analogous to equation 1, with one important exception: While the Q-learning model assumes that the learning rate is constant, in the Kalman filter model, the learning rate is uncertainty-dependent. The trial-wise learning rate *κ_t_* (Kalman gain) depends on the current estimate of the uncertainty of the bandit that is sampled (as per Eq. 8) such that the mean expected value is updated more when bandits with higher uncertainty are sampled. Specifically, ^*σ_ct,t_* refers to the estimated uncertainty of the expected value of the chosen bandit, and *σ_o_* is the observation standard deviation, that is, the variance of the normal distribution from which payouts are drawn (fixed to 4). The uncertainty of the expected value of the chosen bandit is then updated according to

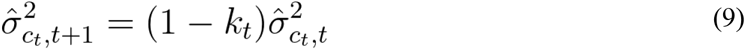

Taken together, this model gives rise to the following intuitions: First, participants not only track the expected mean payoff (*μ*) but also the uncertainty about the expected mean payoff (*σ*). The mean expected value of unsampled bandits is gradually moving towards the decay center and uncertainty about the value increases. Sampling of a bandit leads to a reduction in uncertainty (Eq. 9) that is proportional to the uncertainty prior to sampling. Additionally, the bandit’s mean value is updated via the prediction error (Eq. 7) weighted by the trial-wise learning rate (Eq. 8) such that updating is substantially higher when sampling from uncertain bandits.

We next combined this algorithm for value updating with three different choice rules for action selection. First, we used a standard softmax model (see Eq. 3). Here, choices are only based on the mean value estimates of the bandits *μ_i,t_*, such that exploration occurs in inverse proportion to the softmax parameter *β* and the differences in value estimates:

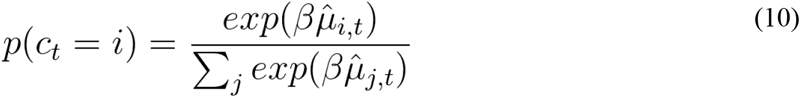

Second, we added an “exploration bonus” parameter *φ* that scales a bandit’s uncertainty ^σ_*i,t*_ and adds this scaled uncertainty as a value bonus for each bandit, as first described by Daw et al. (2006). This term implements directed exploration so that choices are specifically biased towards uncertain bandits.

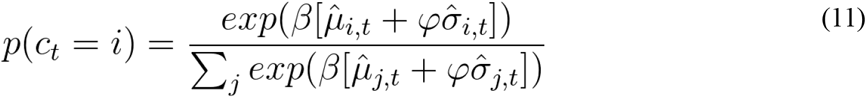

Following a similar logic, we next included a parameter *ρ* modeling choice perseveration. *ρ* models a value bonus for the bandit chosen on the previous trial:

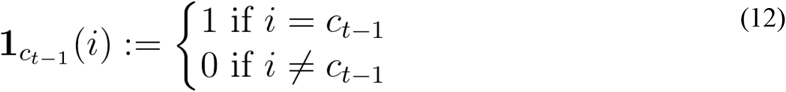

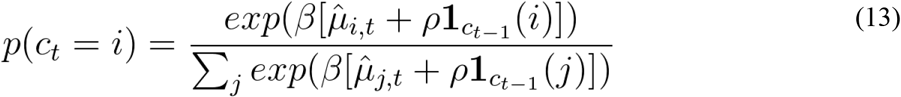

Finally, we set up a full model including both directed exploration (*φ*) and perseveration (*ρ*) terms:

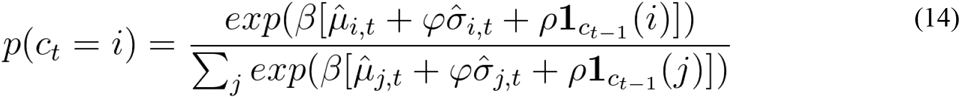

In total, our model space therefore consisted of five models: 1) Q-learning model with softmax, 2) Bayesian learner with softmax, 3) Bayesian learner with softmax and exploration bonus, 4) Bayesian learner with softmax and perseveration bonus and 5) Bayesian learner with softmax, exploration bonus and perseveration bonus. All models were fitted using hierarchical Bayesian parameter estimation in Stan version 2.18.1 (Carpenter et al., 2017) with separate group-level normal distributions for gamblers and controls for each choice parameter (*β*, *φ*, and *ρ*), from which individual-participant parameters were drawn. We ran four chains with 5k warmup samples and retained 10k samples for analysis. Group-level priors for means were set to uniform distributions over sensible ranges (*β*=[0,3]; *φ*=[−20,20]; *ρ*=[−20,20]). Group level priors for variance parameters were set to half-Cauchy with mode 0 and scale 3.

To verify that group differences in the choice parameters (*β*, *φ*, and *ρ*) were not confounded by group differences in the walk parameters, we estimated supplementary models were the random walk parameters were allowed to vary. Parameters were estimated one at a time due to convergence issues, and in a non-hierarchical fashion (i.e. one parameter per group) with the following uniform priors: *λ*=[0,1]; *θ*=[0,100], *σ_d_*=[0,20].

Model comparison was performed using the Watanabe-Akaike Information Criterion, WAIC (Watanabe, 2010; Vehtari et al., 2017) where smaller values indicate a better fit. To examine group differences in the parameters of interest (*β*, *φ*, and *ρ*) we examined the posterior distributions of the group-level parameter means. Specifically, we report mean posterior group differences, standardized effect sizes for group differences and Bayes Factors testing for directional effects (Marsman and Wagenmakers, 2017; Pedersen et al., 2017). Directional Bayes Factors (dBF) were computed as *dBF=i / 1−i* where *i* is the integral of the posterior distribution of the group difference from 0 to +∞, which we estimated via non-parametric density estimation.

### fMRI setup

MRI data were collected with a Siemens Trio 3T system using a 32-channel head coil. Functional MRI (fMRI) was recorded in four blocks. Each volume consisted of 40 slices (2 × 2 × 2 mm in-plane resolution and 1-mm gap, repetition time=2.47s, echo time 26ms). We tilted volumes by 30° from the anterior and posterior commissures connection line to avoid distortions in the frontal cortex (Deichmann et al., 2003). Participants viewed the screen via a head-coil mounted mirror. High-resolution T1 and MT weighted structural images were acquired after functional scanning was completed.

### fMRI preprocessing

MRI data preprocessing and analysis was done using SPM12 (Wellcome Department of Cognitive Neurology, London, United Kingdom). First, volumes were realigned and unwarped to account for head movement and distortion during scanning. Second, slice time correction to the onset of the middle slice was performed to account for the shifted acquisition time of slices within a volume. Third, structural images were co-registered to the functional images. Finally, all images were smoothed (8mm FWHM) and normalized to MNI-space using DARTEL tools and the VBM8 template.

### fMRI analysis

On the first level, we used General Linear Models (GLMs) implemented in SPM12. GLM 1 included the following regressors: 1) trial onset, 2) trial onset modulated by a binary parametric modulator coding whether the trial was a random exploration trial, 3) trial onset modulated by a binary parametric modulator coding whether the trial was a directed exploration trial, 4) outcome onset, 5) outcome onset modulated by model-based prediction error, and 6) outcome onset modulated by model-based expected value of the chosen bandit. Missing responses were modeled separately.

Based on the best-fitting computational model, trials were classified. *Exploitation* trials are trials with choices of the bandit with the highest sum of expected value, uncertainty bonus and perseveration bonus (i.e. the highest softmax probability). *Exploration* trials are all other trials. These were further subdivided into trials on which participants selected the bandit with the highest exploration bonus (*directed exploration* trials) and all other trials (*random exploration* trials). Please note that the trial classification in GLM 1 leads to exploitation trials to be the baseline and exploration as activation relative to this baseline.

GLM 2 included the following regressors: 1) trial onset, 2) outcome onset, and 3) outcome onset modulated by the number of points earned. Missing responses were modeled separately.

GLM 3 is following GLM 1 but replaced the trial classification by the summed uncertainty of all four choice options. Thus, it included the following regressors: 1) trial onset, 2) trial onset modulated by the summed uncertainty of all choice options, 3) outcome onset, 4) outcome onset modulated by model-based prediction error, and 5) outcome onset modulated by the model-based expected value of the chosen bandit. Missing responses were modeled separately.

Group differences were assessed by a second-level random-effects analysis (two-sample t-test). Here we included covariates for depression (BDI-II score), alcohol consumption (AUDIT score), smoking (FTND score), and age. Covariates were z-scored across both groups.

### Dynamic causal modeling

Dynamic causal modeling (DCM, Stephan et al., 2008) is a method to formally test and compare different causal connectivity models underlying the BOLD signal. First, we extracted the BOLD time course of regions of interest (ROIs). Following our previous approach (Chakroun et al., 2020), we defined four ROIs of the right hemisphere based on previous research (Daw et al., 2006; Blanchard and Gershman, 2018): frontal pole (FP), lateral parietal cortex (LPS), anterior insula (aIns) and dorsal anterior cingulate cortex (dAcc, see **Table 2** for coordinates). Time courses were extracted from 5 mm spheres around the single-participant peak within the ROI. See results section for more details on the tested models.

**Table 2:** Region of interest analysis. 10 mm spheres were placed around the coordinates of exploration related activations of previous studies. P-values are small volume corrected. See also (Chakroun et al., 2020, appendix 1 – table 5).

| Reference | Location | Coordinates<br>(x/y/z) | Main effect<br>Directed exploration > exploitation | Controls > Gamblers<br>Directed exploration > exploitation |
| --- | --- | --- | --- | --- |
| (Daw et al., 2006) | R frontal pole | 27, 57, 6 | p=0.012 | No cluster |
| (Daw et al., 2006) | L frontal pole | -28, 48, 4 | p<0.001 | No cluster |
| (Daw et al., 2006) | R lateral parietal cortex | 39, -36, 42 | p<0.001 | p=0.223 |
| (Daw et al., 2006) | L lateral parietal cortex | -29, -33, 45 | p<0.001 | No cluster |
| (Blanchard and Gershman, 2018) | R anterior insula | 32, 22, -8 | p=0.003 | p=0.14 |
| (Blanchard and Gershman, 2018) | L anterior insula | -30, 16, -8 | p=0.002 | p=0.05 |
| (Blanchard and Gershman, 2018) | R dorsal anterior cingulate cortex | 8, 16, 46 | p<0.001 | p=0.048 |

### Classification analysis

To examine whether connectivity dynamics in an exploration-related network was associated with group status (see the previous paragraph), we used an unbiased, leave-one-pair-out approach for group membership classification. We trained a support vector machine classifier (SVM, Chang and Lin, 2011, C=1) on all participants except one patient and one control. The prediction accuracy was computed based on the left-out pair. We repeated this for all possible pairs of controls and gamblers and averaged accuracies across left-out pairs. Finally, we repeated this procedure 500 times with randomly shuffled labels to build a null-distribution, which allows assessing the significance of the observed accuracy.

### Data availability statement

The data that support the findings of this study are available on Zenodo (long format matrix for the behavioral data and t-maps for the fMRI data at, https://doi.org/10.5281/zenodo.4271604).

## Results

### Model-free results

The group difference in the number of points earned was not significant (controls mean[SD]= 18204.83 [1435.37], gamblers mean[SD]= 18489.69 [1520.32], t_43.86_=−0.54, p=0.51). Median response times tended to be shorter in gamblers (controls mean [SD]=0.44s [0.05], gamblers mean [SD]=0.40s [0.07], t_39.413_=1.59, p=0.11). As a model-free measure of exploration, we computed the sequential exploration index (Ligneul, 2019). This index tracks whether in each quadruple of trials all choices are unique, which might reflect a systematic exploration of options. Although this index was numerically higher in controls vs. gamblers (controls mean [SD]=0.065 [0.067], gamblers mean [SD]=0.057 [0.034]), the difference was not statistically significant (t_33_=−0.48, p=0.6).

### Model comparison

Next, we used model comparison based on the Watanabe-Akaike Information Criterion (WAIC, where lower values indicate a better fit) (Gelman et al., 2014) to examine the behavioral data for signatures of directed exploration and perseveration. In both groups, the Bayesian learning model (Kalman Filter) with softmax, exploration bonus, and perseveration bonus accounted for the data best (see **Figure 2**). This model ranking was also observed when the original behavioral data from the Daw et al. (2006) study was re-analyzed using our hierarchical Bayesian estimation approach (see **Figure 2**).

**Figure 2.**
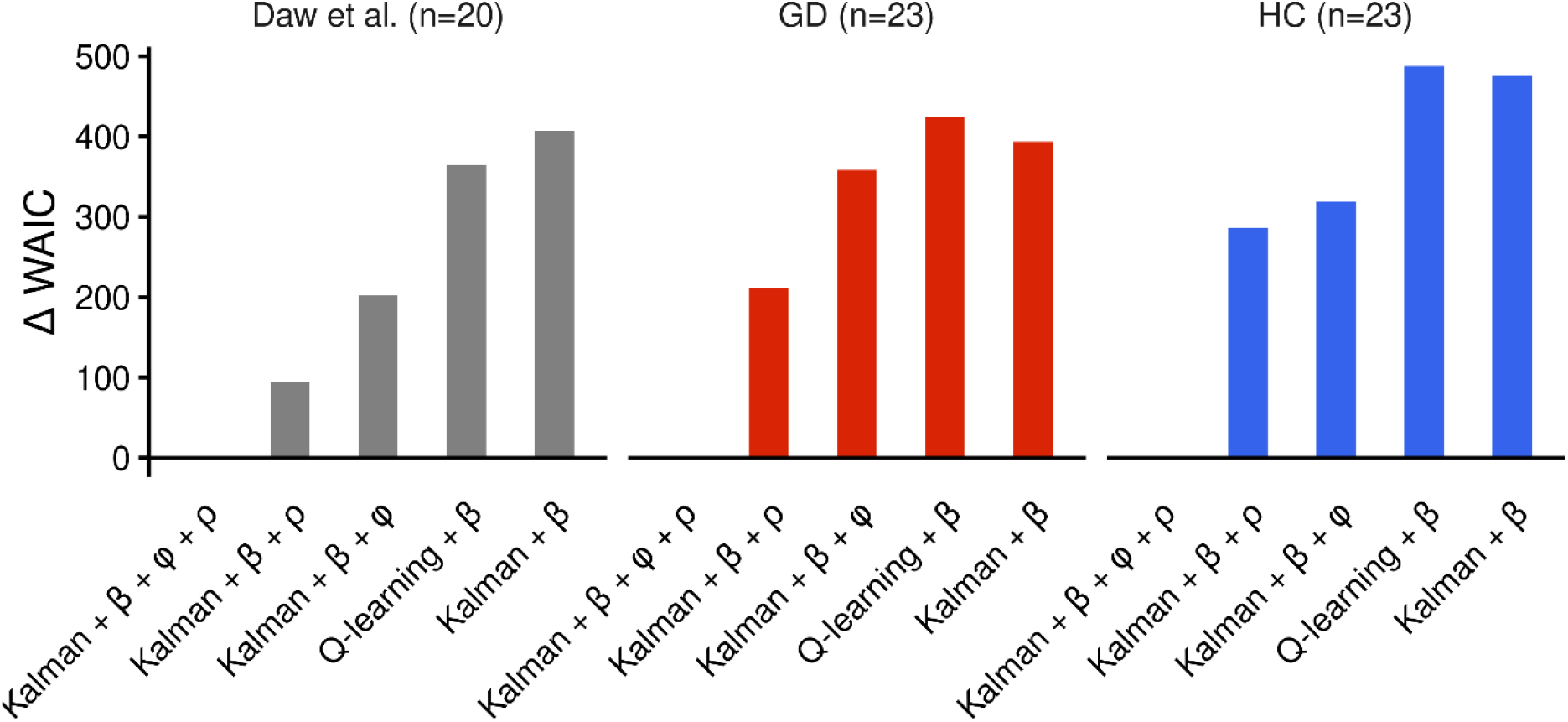
Results of the model comparison based on the Watanabe-Akaike Information Criterion (WAIC). Plotted are WAIC differences between each model and the best-fitting model, such that smaller values indicate a superior fit. Across a re-analysis of behavioral data from Daw et al. (2006) (n=20), gamblers (GD, n=23) and matched healthy controls (HC, n=23) a Kalman filter model with uncertainty bonus (φ) and perseveration bonus (ρ) accounted for the data best.

The best-fitting model gives rise to the following intuitions: First, participants not only track the expected mean payoff (*μ*) but also the uncertainty about the expected mean payoff (*σ*) of the four bandits. The mean expected value of unsampled bandits is gradually moving towards a decay center and uncertainty about the mean value increases. Sampling of a bandit leads to a reduction in uncertainty that is proportional to the uncertainty prior to sampling. Additionally, the bandit’s mean value is updated via a prediction error weighted by a trial-wise learning rate (Kalman gain kappa) such that sampling from uncertain bandits leads to more substantial updating compared to sampling from a bandit with lower uncertainty. Second, action selection is then a function of by the mean expected value of the bandits, an uncertainty bonus (which favors selecting bandits which high uncertainty) and a perseveration bonus (which favors repeating the choice made on the previous trial).

### Parameters of the best-fitting model

Next, we analyzed the parameters of the best-fitting model in detail, focusing on choice stochasticity (softmax slope *β*), exploration bonus (directed, uncertainty-based exploration *φ*), and perseveration bonus (*ρ*, see **Figure 3**). There was evidence for a decrease in *φ* in the gamblers (see **Figure 3 D**), such that a decrease in directed exploration in gamblers was about 12 times more likely than an increase, given the data (dBF = 12.15). Choice stochasticity *β* and perseveration *ρ*, on the other hand, were similar between groups such that the group difference distributions were in each case centered at zero and of inconclusive directionality (dBF = 2.44 and dBF = 1.17, see **Figure 3 B** and **F**).

**Figure 3.**
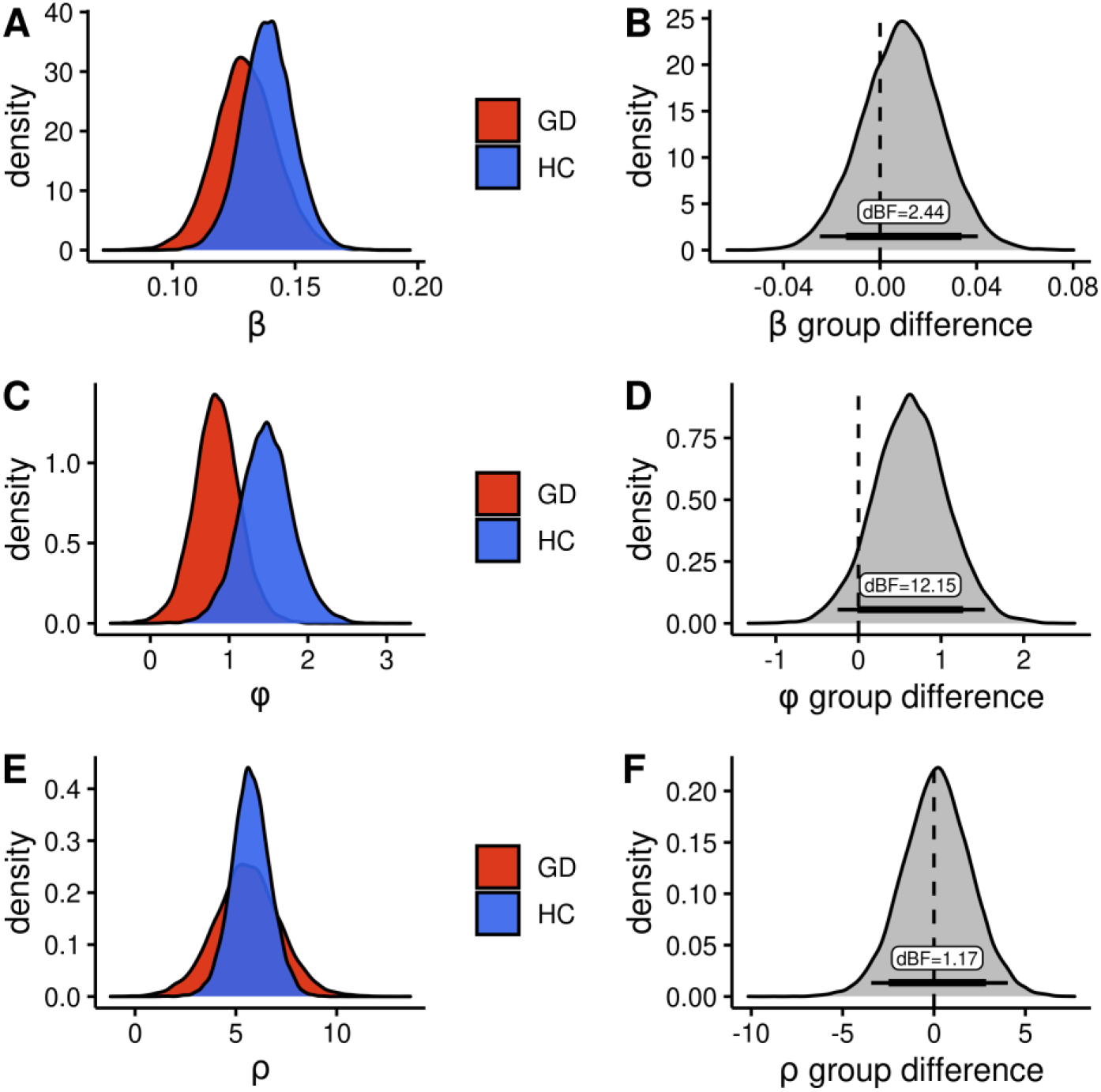
Group parameters of the best-fitting model. GD: Gambling disorder, HC: Healthy controls. **A, C, E:** Posterior distribution of group-level parameters per group. **B, D, F:** Distribution of differences in posterior distributions between groups. Bottom lines indicate the 85% and 95% highest density interval of the distribution. dBF: Directed Bayes factor, the proportion of the difference distribution above 0 over the proportion of the difference distribution below 0. **A** and **B** are showing the β parameter, which represents random exploration. **C** and **D** are showing the φ parameter which represents directed exploration. **E** and **F** are showing the ρ parameter which represents perseveration.

As an additional test of whether groups were statistically distinguishable based on this model, we re-fit the best fitting model with single group-level gaussian distributions per parameter (as opposed to modeling separate group-level gaussians for each group). A model comparison between this model with group-level distributions shared between groups and the original model with separate group-level parameters for controls and gamblers provided further evidence for a group difference: the data were better accounted for by a model with separate vs. shared group-level distributions (WAIC_separate_: 21045.90, WAIC_shared_: 21049.57).

We next explored whether individual differences in gambling addiction severity were associated with exploration behavior in the gamblers. As an index of addiction severity, we computed the mean zscore of SOGS and KFG scores. The correlation between addiction severity and single-participant *φ* parameters was not significant (r=0.01, p=0.95). *φ* parameters also did not correlate with any sub-scale of the Gambling Related Cognition Scale (GRCS, all correlations r<0.2, p>0.38). To explore the effect of potential covariates on *φ*, we performed a regression analysis across both groups. We predicted *φ* based on income, AUDIT-score (alcohol consumption), age, BDI-II (depression), GRCS (gambling cognitions), FTND-score (nicotine addiction), school years and gambling addiction severity. None of the predictors were significant.

### Group differences in random walk parameters

The observed changes in *φ* could have been affected by differences in the representations of the random walks between groups. Because the corresponding parameters (*λ, θ* and *σ_d_*) were fixed in our initial analysis, we ran additional models in which these parameters were free to vary. Due to convergence problems, these parameters were estimated one at a time, and in a non-hierarchical fashion. Still, *σ_d_* could not be estimated reliably due to convergence issues, but allowing *θ* to vary between groups still revealed a group difference in *φ* (dBF = 13.18), and the same was true when *λ* was allowed to vary (dBF = 9.05). While *θ* was similar between groups (dBF =.44), there was evidence for an increase in *λ* in gamblers vs. controls (dBF =.08).

### FMRI

#### Group conjunctions

We first examined standard parametric and categorical contrasts, focusing on conjunction effects testing for consistent effects across groups (GLM2). Ventro-medial prefrontal cortex (vmPFC), ventral striatum (VS) and posterior cingulate cortex (PCC) parametrically tracked outcome value, in line with numerous previous studies and meta-analyses (Daw et al., 2006; Bartra et al., 2013; Clithero and Rangel, 2014, see Figure 4). Importantly, outcome value effects in these regions were observed across controls and gamblers, with no evidence for a group difference.

**Figure 4.**
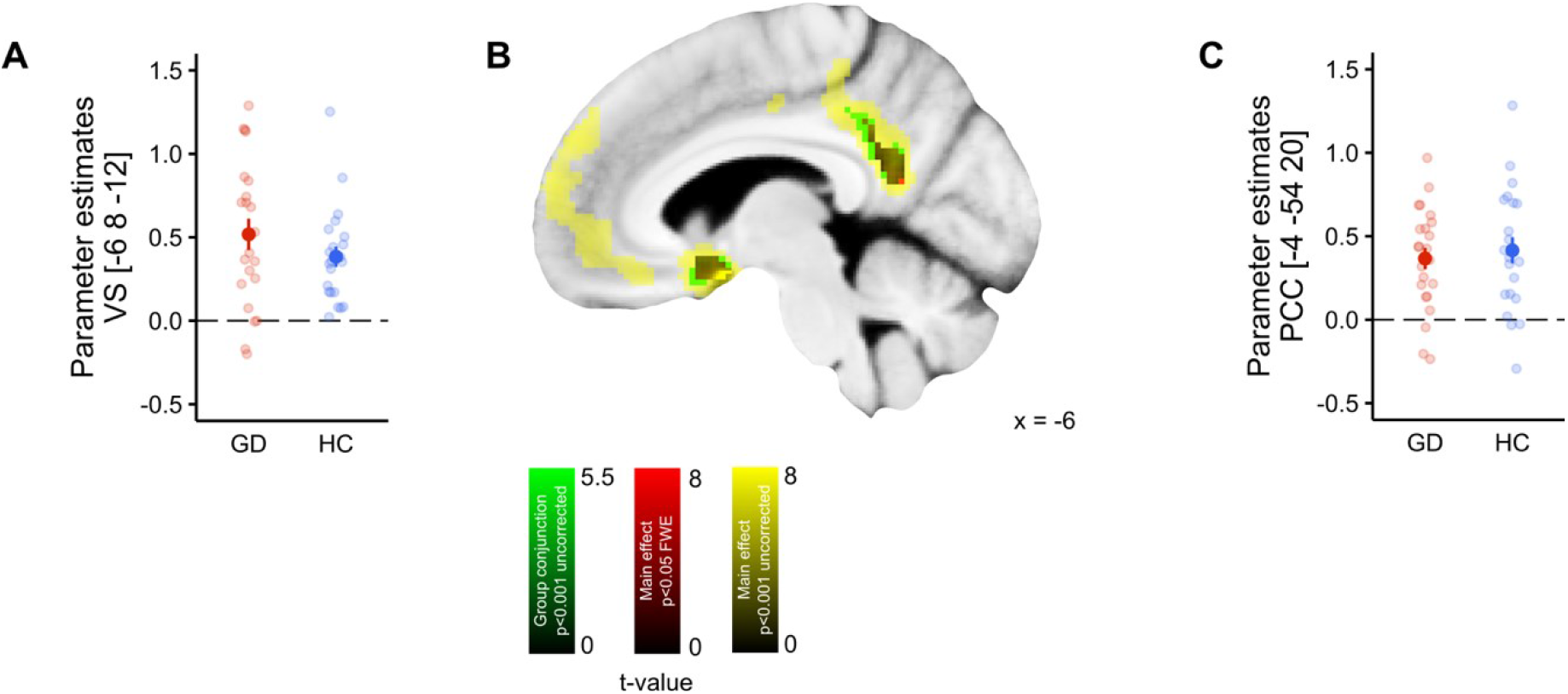
Neuronal correlates of outcome value (points received). A: Extracted beta estimates of each participant in the ventral striatum (VS), peak coordinate of the main effect. **B**: Display of the main effect value, yellow: p<0.001 uncorrected, red: whole-brain FWE corrected p<0.05, green: conjunction GD & HC p<0.001 uncorrected. **C**: Extracted beta estimates of each participant in the posterior cingulate cortex (PCC), peak coordinate of the main effect.

We computed model-based prediction errors for each trial based on the single-participant parameter estimates of the best-fitting model. As previously described in healthy participants (Daw et al., 2006; Pessiglione et al., 2006), the ventral striatum bilaterally coded these prediction errors in both groups (main effect FWE corrected p<0.05: peak at x=−10, y=8, z=−14, z=6.49 and at 16, 10, −14; z=6.11; group conjunction p<0.001 uncorrected: −14, 10, −14, z=4.65; 16, 10, −14, z=4.32).

Based on the computational model, trials were classified as exploitation, directed exploration or random exploration. **Figure 5A and B** show the main effect of directed exploration > exploitation with extensive effects in a fronto-parietal network, replicating previous findings using the same task (Daw et al., 2006; Chakroun et al., 2020). Region-of-interest (ROI) analyses using the same set of ROIs as in our previous study (Chakroun et al. 2020) confirmed significant main effects of directed exploration > exploitation in bilateral frontal pole, bilateral lateral parietal cortex, anterior insula and dorsal anterior cingulate cortex (10mm spheres around the peak coordinates, see **Table 2**).

**Figure 5.**
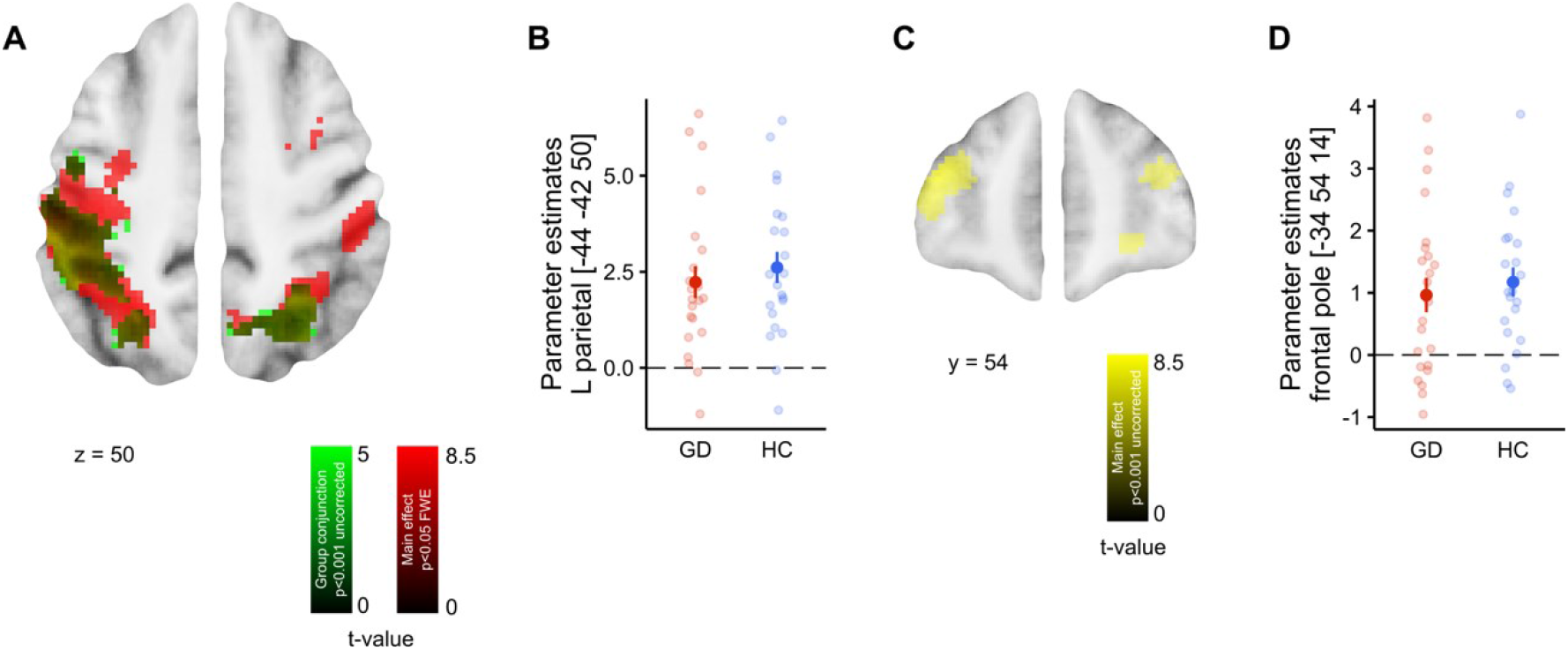
Neuronal correlates of directed exploration. A: Contrast of directed exploration > exploitation in the parietal cortex. red: Main effect whole-brain FWE corrected p<0.05, green: Conjunction HC & GD, p<0.001 (uncorrected). **B**: Parameter estimates per participant from the left parietal cortex. **C**: Contrast of directed exploration > exploitation in the frontal pole, main effect, p<0.001. **D**: Parameter estimates per participant from the left frontal pole.

#### Group differences in exploration-related effects

Next, we next tested our initial hypothesis of reduced frontal pole effects during directed exploration in gamblers (GLM1). We checked for group differences within 10 mm spheres around the peak activations of previous studies (see **Table 2**), which only revealed one group difference in the R dorsal anterior cingulate cortex (p<0.048), which did not survive correction for multiple comparisons across the set of ROIs. We next performed an exploratory whole-brain analysis (at p<0.001 uncorrected) of group differences in brain activity during directed exploration. Controls showed greater activation in parietal cortex (58, −34, 42, z=3.65, p<0.001 uncorrected) and in the substantia nigra / ventral tegmental area (SN/VTA, −12, −18, −10, z=3.78, p<0.001 uncorrected, **Figure 6**). An exploration for effects of gambling severity on exploration-related brain activity in the gambling group revealed no supra-threshold effects even at an uncorrected threshold of p<0.001.

**Figure 6.**
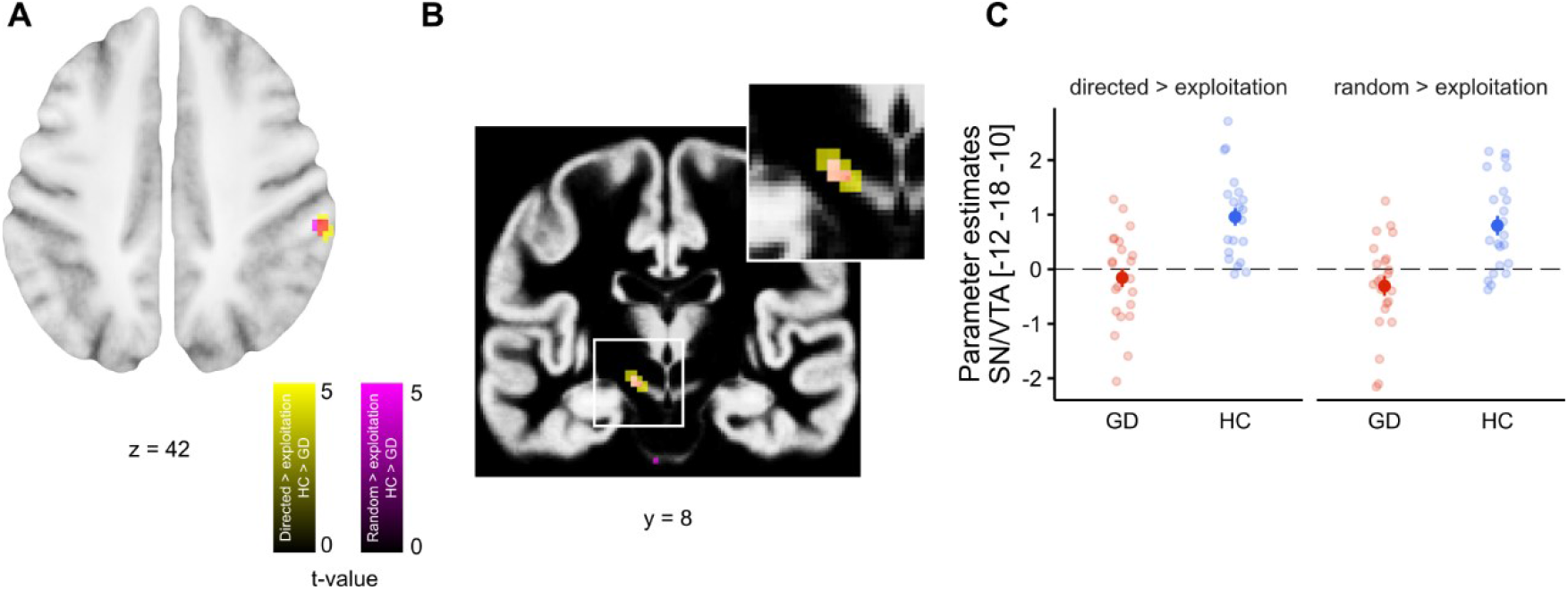
Group differences in the neuronal correlates of directed exploration vs. exploitation. A: Greater activation for controls vs. gamblers in the parietal cortex, p<0.001. **B**: Greater activation for healthy controls compared to gamblers in the SN/VTA, p<0.001. **C**: Parameter estimates per participant from the peak voxel of **B**. Note that the group difference was observed for both directed and random exploration vs. exploitation.

#### Uncertainty-related effects

Following our previous finding that dopamine is involved in the representation of overall uncertainty (Chakroun et al., 2020) and due to the potential involvement of dopamine in group differences, we tested for group differences in the representation of overall uncertainty, that is, the summed uncertainty over all four bandits. In light of our previous findings, we restricted this analysis to regions where we observed these effects before. A direct test at the peak coordinates of Chakroun et al. (2020) with 10mm spherical ROIs revealed a main effect cluster in the dAcc (−3, 21, 39, z= 4.39, pSVC = 0.001), but not in the anterior (42, 15, −6) and posterior (−34, −20, 8) insula. No group differences were observed in these three ROIs.

#### Dynamic causal modeling and group differences in connectivity

To examine whether group differences in network interactions might also contribute to the observed exploration deficit in the gambling group, we used dynamic causal modeling (DCM). For each participant, we extracted the BOLD time-courses in four regions of interest (ROIs) of the right hemisphere based on previous research (Daw et al., 2006; Blanchard and Gershman, 2018): frontal pole (FP), lateral parietal cortex (LPS), anterior insula (aIns) and dorsal anterior cingulate cortex (dAcc, see **Table 2** for coordinates).

As driving input, we used the binary regressor coding directed exploration trials vs. other trials. Because we did not expect structural differences between the groups, all models included all reciprocal connections between the ROIs. We varied the position of the input, ranging from no input to an input to all four ROIs, resulting in 16 models. Bayesian model selection (Stephan et al., 2009) revealed that the model with input confined to the parietal cortex accounted for the data best (expected probability=0.46, exceedance probability=0.99, see **Figure 7B** for a graphical depiction of the best-fitting model). A separate model selection in both groups revealed the same ranking, with model 5 accounting for the data best. Further analysis then proceeded in two steps. First, we extracted single-participant coupling and input-weight parameters (using Bayesian model averaging (Penny et al., 2010) which normalizes extracted parameters by the model evidence per participant). Parameters of the parietal cortex to parietal cortex, frontal pole to parietal cortex and frontal pole to anterior insula connection showed a trend-level difference in gamblers vs. controls: (two-sample t-tests p<0.1, FDR corrected for multiple comparisons, **see Figure 7C**).

**Figure 7.**
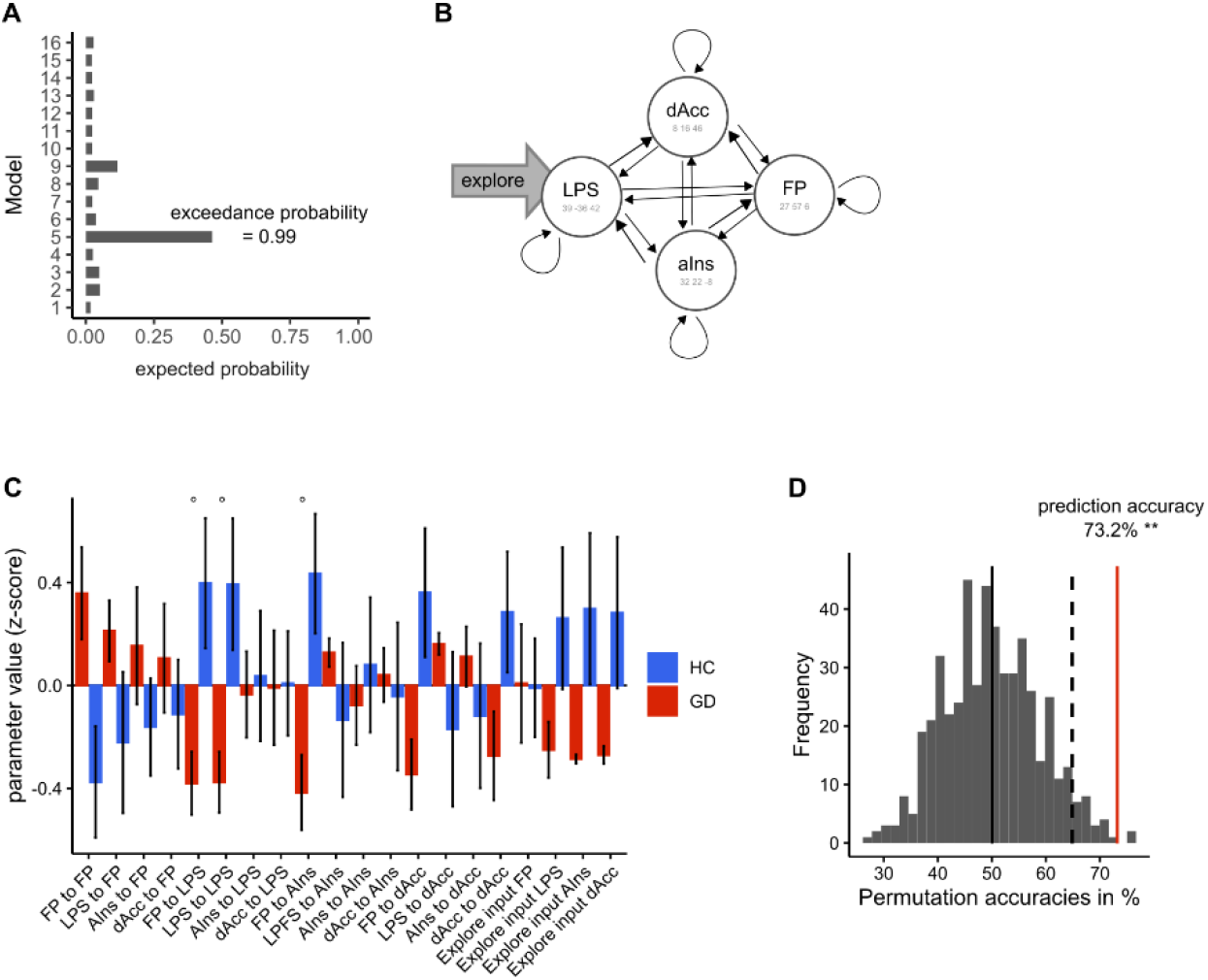
Dynamic causal modeling. A: model selection of 16 models which vary all possible driving inputs. B: Illustration of the DCM winning in the model selection. explore: input of the explore regressor in parietal cortex, LPS: Parietal cortex, AIns: anterior insula, FP: Fronto-polar cortex, dAcc, dorsal anterior cingulate cortex. The gray arrow indicates the driving input. **C**: BMA weighted parameter comparison between groups. GD: Gambling disorder. HC: Healthy controls. ° is indicating trend-level significance (p<0.1, FDR-corrected). **D**: Group identity was predicted based on the DCM parameters (leave-one-pair-out procedure). Statistical significance was assessed with a permutation test. Displayed is the distribution of prediction accuracies under the null hypothesis (500 randomly shuffled labels). The observed accuracy of 73.2% (red line) is beyond the 95% interval of the null-distribution (dashed line).

Second, we tested the hypothesis that the overall connectivity pattern contained information predictive of group (Brodersen et al., 2014). To this end, we used a support vector machine classifier to predict group membership based on all DCM parameters via a leave-one-pair-out, group-size balanced cross-validation scheme. The observed classification accuracy of 73.2% was significantly above chance level (p=0.004, permutation test). Thus, the DCM analyses confirmed that the pattern of functional network interactions contained information about group status, although several univariate analyses in these same ROIs did not reveal group differences.

## Discussion

Here we used a combination of computational modeling and fMRI to investigate exploration behavior in gamblers using a four-armed restless bandit task. Modeling revealed attenuated directed exploration in gamblers. FMRI showed no significant group differences in the representation of basic task variables such as outcome value and reward prediction errors. An exploratory analysis, however, revealed reduced activity during directed exploration in the SN/VTA in gamblers. Dynamic causal modeling showed that coupling in an exploration-related network dissociated gamblers from controls.

In the light of previous findings of reduced behavioral flexibility in gambling disorder, we hypothesized gamblers to show a specific reduction in directed exploration. While both perseveration bonus parameter (*ρ*) and random exploration (*β*) were similar between groups, directed exploration (*φ*) was substantially reduced in gamblers. Estimates of exploration can be confounded by choice perseveration (Payzan-LeNestour and Bossaerts, 2012; Wilson et al., 2014). This is particularly important in gambling disorder where increased perseveration has been reported (van Timmeren et al., 2018). We addressed this issue by extending existing models of exploration with an additional perseveration bonus term, such that final estimates of directed exploration were not confounded by potential group differences in perseveration (see also Chakroun et al., 2020 for a more extensive discussion). Indeed, the full model including both directed exploration and perseveration terms accounted for the data best in both groups. This model ranking was replicated in a re-analysis of the behavioral data from Daw et al. (2006). Notably, our re-analysis of the Daw et al. (2006) behavioral data revealed a contribution of directed exploration that was not observed in their original analysis. As discussed previously (Chakroun et al., 2020), the estimation of directed exploration depends on whether perseveration is explicitly accounted for in the model. Otherwise, the model accounts for perseveration behavior by fitting a reduced (uncertainty-avoiding) exploration bonus parameter.

On the neural level, we found that basic task parameters were similarly represented in both groups. Value effects were localized in a well-characterized network including vmPFC, ventral striatum and posterior cingulate cortex, in line with previous meta-analysis (Bartra et al., 2013; Clithero and Rangel, 2014), with no evidence for group differences. Likewise, striatal prediction error signals were similar between groups. Again, this replicates findings in controls (McClure et al., 2003; Pessiglione et al., 2006). However, the nature of reward signals in gambling addiction remains an issue of considerable debate and inconsistency (Reuter et al., 2005; Balodis et al., 2012; Leyton and Vezina, 2012; Miedl et al., 2012, 2014; Van Holst et al., 2012; Clark et al., 2019). These inconsistencies might be due to specific differences in the implementation and/or analysis of the different tasks. Our version of the four-armed bandit task included neither gambling cues nor monetary reward cues or explicit probability information. These factors may have contributed to the null findings regarding group differences in basic parametric effects of value and prediction error. Few participants in the present sample exhibited very high levels of addiction severity (compared to e.g. Miedl et al. (2012)). This might have precluded us from detecting more pronounced group differences in neural value and prediction error effects. We also did not observe correlations between gambling-related control beliefs and exploration behavior or between addiction severity and behavioral and/or fMRI readouts. While this contrasts with some previous findings using different tasks (Reuter et al., 2005; Miedl et al., 2012; van Holst et al., 2012), overall such effects show considerable variability, both regarding behavior (Wiehler and Peters, 2015) and in reward-related imaging findings (Clark et al., 2019). Our study still included a considerable range of addiction severity (SOGS scores ranged from 3 to 17) suggesting that range restriction is an unlikely explanation for the lack of correlations. However, given the limited sample size typical of studies in such clinical populations, statistical power is an additional concern in the present study.

For the analysis of neural exploration effects, we extended previous approaches (Daw et al., 2006) by separating the neural effects of directed and random exploration via a model-based classification of trials. Again, overall effects were highly similar between groups and consistent with previous studies, such that directed exploration recruited a fronto-parietal network including frontal pole regions (Daw et al., 2006; Badre et al., 2012; Raja Beharelle et al., 2015; Chakroun et al., 2020). Importantly, our initial hypothesis of attenuated frontal pole activation in gamblers was not confirmed. Although frontal pole and IPS effects of directed exploration were numerically smaller in gamblers (see Figure 6), neither group difference was significant. However, the DCM analysis revealed was some evidence for altered frontal pole connectivity in gamblers (i.e. trend-level reductions in connectivity for frontal pole-parietal cortex and frontal pole-anterior insula interactions, see Figure 7C).

Given that dopamine has been implicated in both the exploration/exploitation trade-off (Frank et al., 2009; Beeler, 2012; Kayser et al., 2014; Gardner et al., 2018; Chakroun et al., 2020) and gambling disorder (Voon et al., 2006; Boileau et al., 2014; Majuri et al., 2017; van Holst et al., 2017; Mathar et al., 2018; Potenza, 2018), we additionally carried out an exploratory analysis of subcortical correlates of directed exploration. The finding of reduced SN/VTA effects during exploration in gamblers resonates with a recent study that reported increased dopamine synthesis capacity in striatal regions in gamblers (van Holst et al., 2017), but see Potenza (2018) for a critical discussion. Given the reciprocal connectivity between striatum and SN/VTA (Haber and Knutson, 2010), increased striatal-midbrain feedback inhibition might be one mechanism underlying this effect. However, small midbrain effects can be affected by cardiac or respiratory artifacts, which were not directly controlled in the present study.

Group membership could be decoded from the DCM coupling parameters with an accuracy of 73.2%. This supports the idea that network interactions might contain additional information reflecting a participants’ clinical status compared to univariate contrasts (Brodersen et al., 2014). However, we emphasize that the overall prediction accuracy, although significantly above chance-level based on permutation testing, is still too low for potential clinical applications. Higher accuracies could be achieved by larger training data sets (to reduce the noise induced by individual outliers). Also, more detailed network models might better reflect the underlying neural computations and thus yield higher predictive accuracy.

A number of limitations of the present study need to be acknowledged. First, it remains unclear whether reduced directed exploration constitutes a vulnerability factor or a consequence of continuous gambling. It would therefore be interesting to see whether these effects are tied to the clinical development of patients (e.g. to the escalation of gambling behavior or treatment effects) or whether they manifest as stable factors that increase the risk for the development of the disorder. Second, a comparison of the present results to patients with substance-use disorders would be of considerable interest (Morris et al., 2016), in particular given the overlap in terms of decision-making impairments. Third, tasks such as the observe-or-bet task (Blanchard and Gershman, 2018) or the Horizon Task (Wilson et al., 2014) might allow for a more clear-cut dissociation between directed and random exploration than the bandit task employed here. The reason is that the computational model assumes that value, perseveration and exploration bonus jointly affect action probabilities at the time of choice. Our classification of trials into exploitation, directed exploration and random exploration based on the fitted model therefore might not constitute as clear-cut segregation of the involved processes as in these other tasks. On the other hand, one advantage of the bandit task is that it assesses behavior as it unfolds over longer learning periods in a dynamic environment. As such, it might better resemble exploration behavior as it occurs in dynamic real-world settings. Finally, models with free random walk parameters showed convergence problems, such that these parameters (with the exception of *σ_d_*) could only be estimated in a non-hierarchical fashion (Raja Beharelle et al., 2015; Chakroun et al., 2020). Allowing *λ* or *θ* to vary still revealed a group difference in *φ*. Likely more data is required to reliably estimate these parameters, in particular in clinical samples such as the present one.

Impairments in reward-based learning, decision-making and cognitive control are hallmarks of gambling disorder (Wiehler and Peters, 2015; Clark et al., 2019). Here we show using computational modeling that during reinforcement learning in volatile environments, gamblers’ behavior is characterized by attenuated directed exploration rather than increased perseveration. Whether alterations in the exploration/exploitations trade-off extend to other tasks or environmental statistics, or could account for previous findings of reversal learning impairments in gamblers (de Ruiter et al., 2009; Boog et al., 2014) are interesting open question. Coupling parameters from a dynamic causal model of an exploration-related network contained information predictive of clinical status, raising the possibility that such network interactions might be more diagnostic of this disorder compared to univariate effects. An exploratory analysis of subcortical exploration-related group differences revealed reduced activity in the SN/VTA in gamblers, complementing accumulating evidence for dopaminergic dysfunction associated with this disorder (Boileau et al., 2014; van Holst et al., 2017; van Timmeren et al., 2018; Clark et al., 2019; Kayser, 2019). Taken together, our findings highlight computational mechanisms underlying reinforcement learning in volatile environments in gambling disorder. In light of earlier results (Chakroun et al. 2020) this might be related to dopaminergic dysregulation.

## Acknowledgments

A.W. and J.P. designed research. A.W. performed research. A.W. and K.C. analyzed the data. A.W. and J.P. co-wrote the paper, and K.C. provided revisions. J.P. supervised the project. We thank Anica Bäuning for assistance with task programming. We thank Raymond Dolan and Nathaniel Daw for kindly making the behavioral data from their 2006 paper available for re-analysis.

